# DNA Extraction Protocols for Animal Fecal Material on Blood Spot Cards

**DOI:** 10.1101/2024.11.03.621776

**Authors:** Ann-Katrin Llarena, Thomas H.A. Haverkamp, Wenche Støldal Gulliksen, Kristin Herstad, Arne Holst-Jensen, Eystein Skjerve, Lisbeth Rannem, Sabrina Rodriguez-Campos, Øivind Øines

## Abstract

**Background:** Collecting fecal samples using dry preservatives is an attractive option in large epidemiological studies as they are easy to use, cheap and independent of cold chain logistics. Here, we test four DNA extraction methods with the aim of identifying an efficient procedure to extract high-quality DNA from fecal material of canine, sheep, equine, bovine, and pig collected on dry blood spot cards, with the goal of generating good quality shotgun metagenomics datasets. Further, the suitability of Illumina shotgun metagenomic sequencing at 20million PE read depth was assessed on its ability to successfully characterize the taxonomic and functional aspects of the resulting fecal microbiome.

**Methods:** DNA was extracted from pig feces and mock communities collected on blood spot cards using four DNA extraction methods; two different methods of the QIAsymphony® PowerFecal® Pro DNA Kit, the ZymoBIOMICS™ DNA Miniprep Kit, and the MagNA Pure 96 DNA and Viral NA Small Volume Kit. Possible extraction bias was controlled by amplicon sequencing of mock communities. Fecal samples from canine, sheep, equine, bovine, and pig were thereafter subjected to the best performing DNA extraction method and shotgun metagenomic sequencing to determine sequencing efforts for functional and taxonomic analysis.

**Results:** The four DNA extraction methods demonstrated similar community composition in the sequenced bacterial mock community. The QIAsymphony® PowerFecal® Pro DNA Kit was identified as the DNA extraction method of choice, and the resulting DNA was subjected to shotgun metagenomic sequencing with 20million PE reads. We found that higher number of reads increased the richness of observed genera between 100,000 and 5 million reads, after which higher sequencing effort did not increase the richness of the metagenomes. As for functional analysis, the number of low abundance functions in the metagenomes of the animals’ feces increased with sequencing depth above 20 million PE reads.

**Conclusion:** Our experiments identified several methods suitable for extracting DNA from feces collected on blood spot cards. The Qiagen’s Blood and Tissue kit on the QiaSymphony platform fulfilled the criteria of high yield, quality, and unbiased DNA, while maintaining high throughput for shotgun metagenomic sequencing. A sequencing depth of 20 million PE reads proved adequate for taxonomic estimations and identifying common functional pathways. Detecting rarer traits, however, requires more sequencing effort.

## Introduction

The microbial community in the gastrointestinal tract plays an important part in physiology and health of animals and humans. The gut microbes closely interact with the host’s epithelial cells and are important for digestion, immune response, neuro-gastro pathways, and development of metabolic and inflammatory illnesses (1–3). The intestinal tract can also harbor numerous animal and human pathogens and antimicrobial resistance (AMR) genes and bacteria transmissible to humans, animals, and the environment (4–7), constituting a possible public health risk. Therefore, the microbial community in the animal gut has been under scrutiny in numerous studies (8–14).

Using feces as a proxy for gastrointestinal luminal content is a simple and non-invasive method to collect samples and animal owners can easily collect feces. Feces is discharged to the environment, and animal and human pathogens with a fecal-oral route of transmission makes feces relevant to study in a One Health context (7). However, different collection methods, transport, and storage time and conditions can affect sample integrity (15–19), and thus the inference of the microbial community composition and nucleic acids profile, albeit the variation observed among individuals is expected to exceed the bias induced by various methods and approaches for sampling (17, 19, 20). Sampling, transport, and storage procedures should nevertheless be selected to minimize post-collection bias in microbial community composition. The presumptive gold standard is considered to be rapid freezing (20, 21), or the use of liquid preservatives such as 95 percent ethanol, DNA/RNA Shield (Zymo Research), and RNAlater (ThermoFisher). These methods are impractical in large-scale field research where the collection method must be simple, independent of cold chain logistics, and often have budget constraints (22). Dry collection methods involve using containers, material or devices that preserve the fecal samples in a dry state. These can be dried blood spot (DBS) cards made of protein-binding cellulose, which have been in use in diagnostics since 20^th^ century (23), and more recent materials made of papers treated with chemicals to lyse microorganisms and bind proteins and nucleic acids, such as occult blood test cards (FOBT) and Flinders Technology Associates (FTA) cards (Whatman PLC, UK). These cards are designed to keep the sample dry and stable during transportation and storage and are appealing choices for sampling fecal material in large field investigations since they are simple to use, easily transportable, can be kept at room temperature, and do not require any additional preservative chemicals. Dry preservatives preserve human feces efficiently (18, 19, 24), including during long term storage (25, 26), but they allow less sample material to be collected and makes it impossible to measure its initial weight (as reviewed by (27)). A few studies have used dry cards to collect fecal material from primates (28), birds (29), and cats (30) intended for microbiota profiling using amplicon sequencing, finding that dry preservatives preserved DNA, albeit with low yield. This limitation must be considered when choosing library protocols, sequencing technique, and choice of bioinformatics (30).

The Trøndelag Health study (HUNT) One Health is a large cohort study that collected feces and health data from approximately 3000 animals (canine, sheep, equine, bovine, and pig) (31). The animal owners collected the fecal samples on DBS cards, air-dried the samples, and sent them to a laboratory for storage. The goal was to perform DNA extraction and shotgun metagenomics sequencing to offer metagenomic datasets suitable for functional and taxonomic characterization to the greater research community. Adequate yield and high-quality DNA are required, as low DNA levels can result in higher biases in GC content, k-mers, or target (e.g. 16S rRNA) gene compositions (32). A critical lower amount of environmental DNA suitable for shotgun metagenomics has been reported to be 10pg of DNA (32), and therefore the selected DNA extraction method for animal fecal material on DBS should yield at least this amount. To date, no reports have clearly demonstrated what is the most suitable method for extracting DNA from animal feces collected on DBS cards. Furthermore, whereas research on the taxonomic and functional characterization of fecal metagenomes using amplicon sequencing are available for all relevant species, few studies use shotgun metagenomics to characterize the fecal metagenome from livestock animals (as reviewed in (33)).

The main aim of the present study was to identify a suitable and common procedure for high-throughput acquisition of shotgun metagenome data from dried animal feces collected on DBS cards. For this, the chosen extraction method should achieve sufficient yield and high-quality DNA, minimized bias in metagenome DNA profiles, and provide sufficient data to allow for the desired level of characterization of taxonomic and functional properties present in the fecal microbiome(s). The chosen procedure was to be successively applied to all animal feces samples collected in the HUNT One Health study.

## Materials and Methods

### Materials

#### Animal feces on DBS cards

Fecal samples were collected after natural defecation in HUNT One Health. The samples were collected by untrained animal owners, based on instructions on paper and video. The owners smeared animal fecal material on two sampling fields of 1.7 x 3.5cm DBS filter paper pieces contained in a customized wrap-around card (Lipidx, Oslo, Norway), hereafter referred to as “DBS card” (Fig 1). We therefore used the same type of DBS filter paper to allow simulation of samples in the HUNT One Health project. Fecal samples of healthy canines, sheep, equines, bovines, and pigs were collected and smeared as a thin layer on the DBS cards, dried for at least two hours and stored at −20°C before pieces of this material was subjected to DNA extraction.

**Fig 1.**
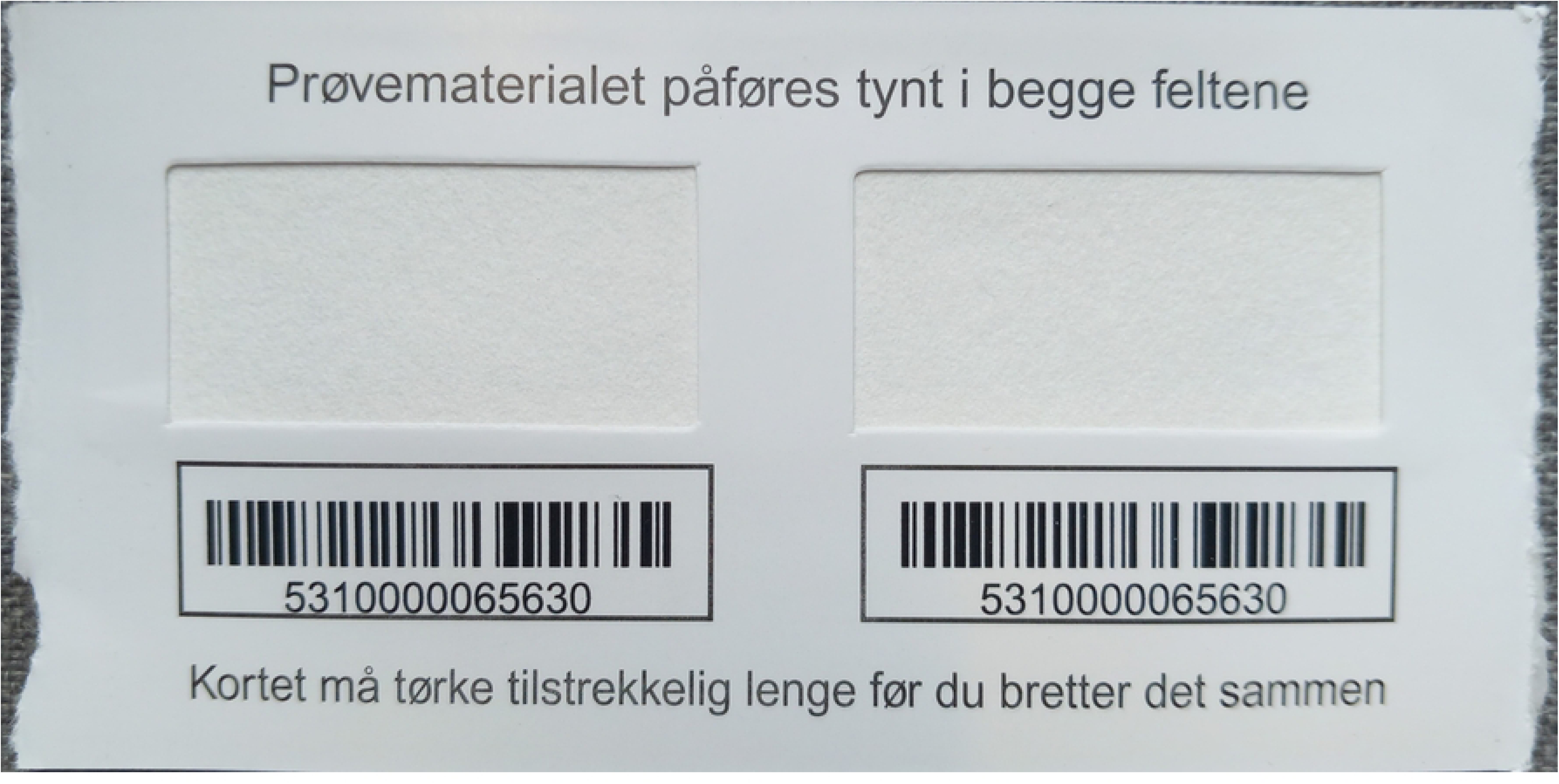
Picture of an empty DBS card with two fields of 170mm*350mm each, equaling an area of ∼12cm2 available for sampling.

Upon extraction, the DBS cards were taken out from the freezer and thawed. Four 8-mm diameter circles were aseptically clipped from each DFC with a single-use biopsy puncher (cat number 273693, Kruuse Norway, Drøbak, Norway). The operators visually ensured similar and representative sections of the DBS with evenly spread fecal material to be selected for sampling. The paper circles were transferred to PowerBead Pro Tubes (2 ml) (cat. no. 19301, Qiagen) containing pre-loaded homogenization beads.

Sampling of feces from animals does not require institutional animal care and use committee (IACUC) approval according to Norwegian legislation (Forskrift om bruk av dyr i forsøk) which is in line with European legislation (Directive 2010/63/EU of the European Parliament and of the Council, Article 40). All owners signed an informed consent form before enrolment.

#### Positive and negative controls

To prepare positive and negative controls, four 8-mm diameter paper discs were aseptically excised from a clean DBS and deposited in a PowerBead Pro tube. Positive controls were spiked with 75 µl of the standard microbial community II from Zymobiomics (Cat. No. D6310, Zymo Research Corporation, USA), hereafter referred to as “mock”. This mock community consists of known quantities of Gram-negative and Gram-positive bacteria together with yeast with varying sizes and cell wall composition. The theoretical composition in terms of 16S rRNA gene abundance as given by the producer, calculated from theoretical genomic DNA composition with the following formula: 16S rRNA gene copy number = total genomic DNA (g) × unit conversion constant (bp/g) / genome size (bp) × 16S copy number per genome, is 95.9% *Listeria monocytogenes*, 2.8% *Pseudomonas aeruginosa*, 1.2% *Bacillus subtilis*, 0.069% *Escherichia coli*, 0.07% *Salmonella enterica*, 0.012% *Lactobacillus fermentum,* 0.00089% *Enterococcus faecalis*, and 0.000089% *Staphylococcus aureus* (Zymobiotics Research Corpooration, USA). Negative controls were DBS paper only, subjected to the same buffers and procedures as the samples containing fecal material, and is hereafter referred to as “blank”.

#### Experimental set up and DNA extraction methods

##### Determine an appropriate DNA extraction method for DBS

To find a suitable DNA extraction method, four different extraction procedures were assessed based on their ability to produce high yield and pure unbiased DNA from pig fecal material and the mock community on DBS cards. Further, to compare how efficiently the selected method extracted DNA from feces from the animal species included in the HUNT One Health project, canine, sheep, equine, bovine, and pig fecal material collected on DBS cards were subjected to DNA extraction (Fig 2).

**Fig 2.**
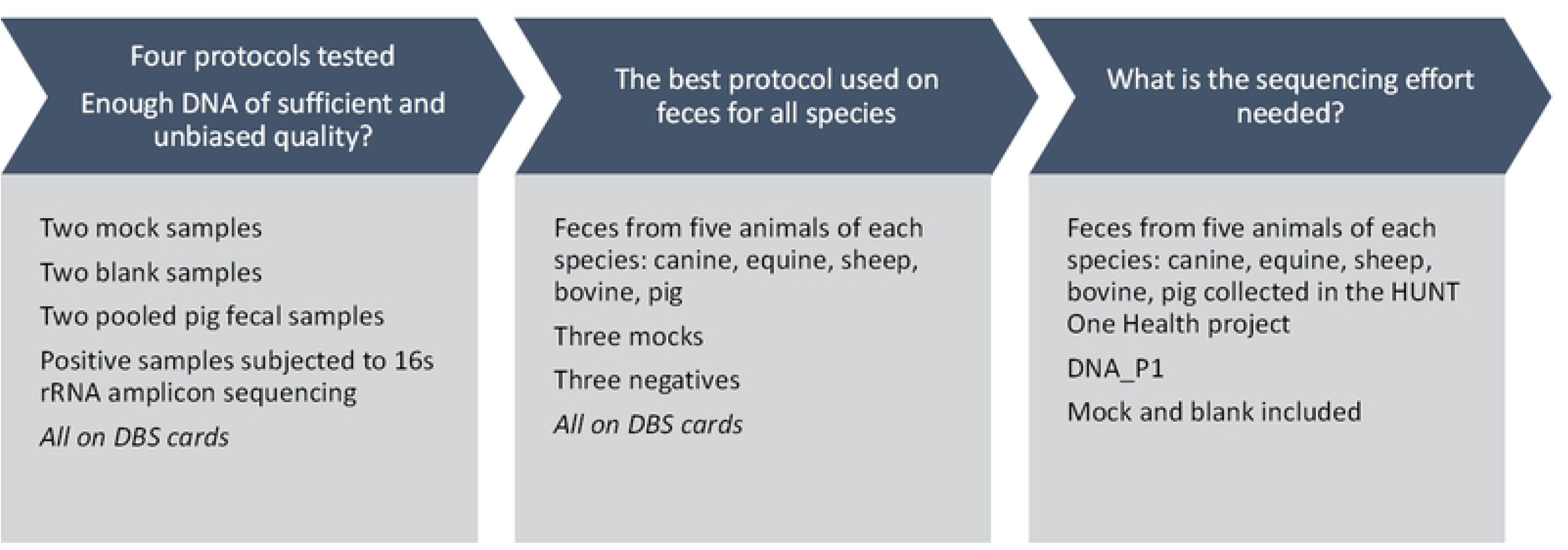
Experimental set up to identify a suitable DNA extraction and purification protocol for DBS. The different methods tested were DNA_P1) QIAsymphony® PowerFecal® Pro DNA Kit (cat. no. 938036, Qiagen) including six rounds of homogenization on a FastPrep-24™ Classic at 6 m/s for 60 seconds, separated with 5 minutes of resting, followed by digestion with proteinase K 600 mAU/ml (Qiagen) prior to automated purification on the QIAsymphony® SP robot (Qiagen). DNA_P2) Performed identical as DNA_P1, but with no proteinase K treatment. DNA_P3) ZymoBIOMICS™ DNA Miniprep Kit (Zymo Research Corp., Irvine, CA, USA) after homogenization using a FastPrep-24™ Classic at 6m/s for 60s for six rounds interspaced with 5 min resting. DNA_P4) MagNA Pure 96 DNA and Viral NA Small Volume Kit on the MagNA Pure 96 Instrument (Roche, Basel, Switzerland) with proteinase K pre-treatment step and standard buffers using the DNA Blood ds SV Protocol optimized for double stranded DNA and Next Generation Sequencing. Of these, DNA_P1, DNA_P2, DNA_P3 and DNA_P4 were used to extract DNA from mocks, blanks and pig feces, DNA_P1 was tested on their performance on fecal from all studied animal in addition to triplicates of positive and blank controls. DNA was extracted from bovine, equine, canine, sheep, and pig feces using DNA_P1 and sequenced on an Illumina NovaSeq to generate >20million PE reads from each sample. These datasets were used to inform on required sequencing effort.

Pig fecal material was chosen as initial representative for the five species as it was found to be challenging to extract sufficient DNA from this matrix in an early pilot (data not shown). The pig feces were collected from two healthy adult pigs, pooled, and properly mixed before being smeared as a thin coating on DBS cards and used in all four protocols. The following DNA extraction methods were tested: DNA protocol 1 (DNA_P1): QIAsymphony® PowerFecal® Pro DNA Kit (cat. no. 938036, Qiagen) including six rounds of homogenization using the FastPrep-24™ Classic (MP Biomedicals, Irvine CA, USA) at 6 m/s for 60 seconds, separated with 5 minutes of resting, followed by digestion with proteinase K 600 mAU/ml (Qiagen) prior to automated purification on the QIAsymphony® SP robot (Qiagen). DNA_P2 was performed identical as DNA_P1, but with no proteinase K treatment. DNA_P3 was ZymoBIOMICS™ DNA Miniprep Kit (Zymo Research Corp., Irvine, CA, USA) after homogenization using a FastPrep-24™ Classic at 6m/s for 60s for six rounds interspaced with 5 min resting. DNA_P4 was MagNA Pure 96 DNA and Viral NA Small Volume Kit on the MagNA Pure 96 Instrument (Roche, Basel, Switzerland) with proteinase K pre-treatment step and standard buffers using the DNA Blood ds SV Protocol optimized for double stranded DNA and Next Generation Sequencing (Table 1). Details on the DNA extraction methods can be found in Supplementary materials. The obtained elute purity and yield was assessed, as well as its capability for high throughput, and the resulting DNA from the mock communities was subjected to amplicon sequencing targeting the bacterial 16S rRNA gene using the Oxford Nanopore MinION device (see chapter “Sequencing methods”).

**Table 1.**
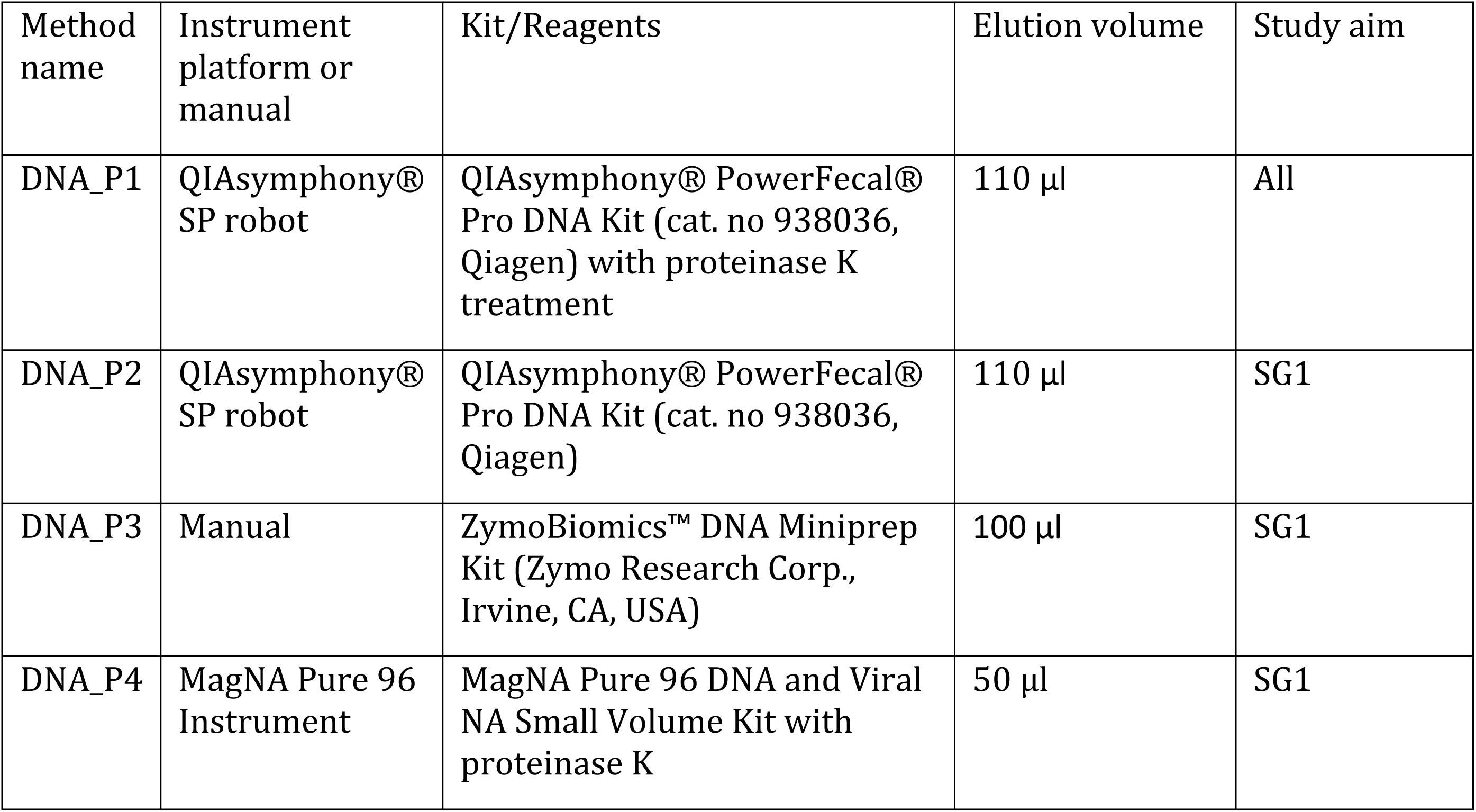
Overview of extraction methods tested.

The best performing protocol considering DNA yield, purity, applicability for high-throughput processing, and ability for generating unbiased sequence data, was used to extract DNA from fecal samples collected from five individuals of each species: canine, sheep, equine, bovine, and pig (in total n=25, Fig 2). Triplicates of mock and blank controls were included in each protocol (Fig 2). The DNA purity and yield was measured and its combined potential for high throughput processing was assessed.

**The DNA concentration was measured** fluorometrically by Qubit™ (Qubit, Thermo-Fisher Scientific, Waltham MA, USA) using the Qubit dsDNA BR (Broad Range) Assay Kit or, if the concentration was too low to yield a result, the Qubit™ dsDNA HS (High Sensitivity) Assay Kit. DNA integrity was assessed by agarose gel electrophoresis either in house or by the sequencing provider, while DNA purity was assessed using a Nanodrop Spectrophotometer (Thermo-Fisher Scientific, Waltham MA, USA) at absorbance wavelength ratio 250nm/230nm against 260nm. A ratio > 1.8 was considered pure. Descriptive statistics for each species (n=2 for pig in first round of experiments, n=5 for all species to test the best method) and extraction method (n=4) were calculated in STATA v.15.1 SE for Windows (StataCORP LLC, Texax, USA) and visualized using Microsoft Office 2016 Win 32 Excel (Microsoft, California, USA).

**To determine sufficient sequencing efforts needed** for functional and taxonomical characterization of the fecal metagenomes of canines, sheep, equines, bovines, and pigs, fecal samples collected on DBS cards of said species were subjected to DNA extraction and shotgun metagenomic sequencing (Fig 2). Feces from five individuals of each species were collected on DBS between 2017 and 2019 in the HUNT One Health Project and extracted using DNA_P1.

Together with a mock community (described above) and a blank sample, the DNA was submitted to sequencing (∼20 million PE reads) via library preparation using the ThruPLEX library preparation kit (Takara, Shiga, Japan), and 150 bp PE sequencing on the Illumina NovaSeq platform (see chapter “Sequencing strategies”).

##### Library preparation and sequencing strategies

DNA extracted from mock samples using DNA_P1 through DNA_P4 was subjected to 16S amplicons sequencing using the nanopore sequencing device MinIon (Oxford Nanopore Technologies, Oxford, UK). The amplicons were generated from the DNA-templates with a PCR using the universal primers 27F/S-D-Bact-0008-c-S-20 and 1492R/S-D-Bact-1492-a-A-22 for long read characterization of bacterial communities with a long-read NGS protocol (34) using a recombinant DNA polymerase (EP0402, ThermoFisher Scientific, Waltham, USA). Amplicons were purified using Macherel-Nagel Nucleospin Gel and PCR cleanup-kit (Düren, Germany), quantified with Qubit 1x HS DNA assay (ThermoFisher Scientific), before up to 440 ng of purified amplicons were subjected to nanopore library preparations using the rapid barcoding kit (SQK RBK004, v RBK_9054_v2_revM_14Aug2019) according to manufacturer’s description with one modification: double volumes were applied to allow top up of the flow-cell (FLO-106D) after a period of sequencing. In total, the Minion flow-cell ran for 28 hrs.

To obtain FASTQ files, the fast5 data was base called using Guppy (version 6.5.7) with default settings (Oxford Nanopore). The output file from Guppy was used for quality control with Nanoplot (version 1.33.1 (35)). A total of 2.4 x 10^6^ reads were produced with a median quality Phred score of 12. The fastq reads were split on adapter using the script duplex_tools (ONT) to generate simplex reads. All FASTQ files were concatenated and subsequently demultiplexed with qcat (version 1.1.0; https://github.com/nanoporetech/qcat) to generate a single FASTQ file per sample with the following settings: minimum read length 50 bp, trim adapters and barcode sequences, and detect-middle. Next, we used NanoFilt (version 2.8.0) to select only reads with an average Phred quality score of 9 or higher with a read length between 1450 and 1650 bp, matching the size of the 16s rRNA (35). Classification of the estimated number of reads based on relative abundance was performed with the tool EMU, using a minimum abundance of 0.0001 with the EMU 16s rRNA classification database as well as the SILVA database (36). The EMU results were imported into R-studio (version 2022.07.01) to visualize the diversity of the samples. The following packages were used in R-version 4.2.1: tidyverse (v1.3.2); ggplot2 (v3.5.1); microViz (v0.12.4)(37).

Library preparation and whole genome Illumina shotgun metagenomic sequencing was performed by BGI Tech Solutions (Hong Kong, China). Libraries were prepared using the ThruPLEX® DNA-seq by Takara (cat. no. R400674, Takara BioInc, Europe), which can generate libraries from as little as 50 pg of DNA. The concentration and fragment length of the resulting libraries were quality checked on a 2100 Bioanalyzer (Agilent, Santa Clara, US) with an in-house qPCR to meet the criteria of Illumina Novaseq sequencing requirements. The libraries were thereafter sequenced on the Novaseq 6000 using the 150 bp PE sequencing strategy, delivering at least 5Gb data per sample.

Quality control and classification of Illumina shotgun metagenomic datasets was first performed by the external sequencing provider (BGI), which delivered clean reads, from which adaptor sequences, contamination and low-quality reads had been removed using SOAPnuke software (version 2.1.0)(38). In brief, SOAPnuke removed the entire read if it contained >25% adapter sequence, >50% bases with a quality score < 20, or >3% N bases. Next, duplicate reads with identical bases were removed from the data. This clean sequence data was processed using the Nextflow pipeline TALOS (https://github.com/NorwegianVeterinaryInstitute/Talos). In brief, the pipeline filters low-quality and low complexity reads and reads matching the human/phix (NC_001422) genomes (39). The cleaned data was then subsampled with the seqtk; version 1.3) into datasets with 10k, 50k, 100k, 500k, 1Million, 5M, and 10M PE reads. Each dataset was then classified taxonomically using Kraken 2 (40), and the PlusPFP database (date: 12/9/2022; https://benlangmead.github.io/aws-indexes/k2). In addition, functional classification with SUPER-FOCUS (41) using the Diamond version 1 database with cluster size 100 (https://github.com/metageni/SUPER-FOCUS#downloading-prebuilt-databases) was performed (42). To identify compositional differences between host species, Sourmash (43) was used to generate kmer sequence signatures from each sample. The signatures were used to generate a Jaccard distance matrix and were visualized using Sourmash plot. The taxonomic and functional classification results were imported into R studio (version 2022.07.1) running R (version 4.2.1). The r-package tidyverse (1.3.2) was used for reformatting the data and visualization of the results. Vegan (2.6-4) (44) was used for calculation of alpha-diversity metrics for both the Taxonomic and functional classifications.

##### Data availability

The Nanopore reads are available under project number PRJEB80558, and the shotgun metagenomes generated by Illumina sequencing are available under project number PRJEB80559.

## Results

To identify a suitable DNA-extraction method for the fecal material collected in the HUNT One Health project, the combined results from DNA concentration measurements, efficiency, microbial community biases introduced, and level of automation and cost of different extraction methods, were compared.

### DNA concentration and yield

The median DNA concentration from mock and pooled pig fecals was 0,34 (range 0,1 – 0,88) ng/µl and 26,8 (range 7,9 – 75,6) ng/µl, respectively. Blanks were always below detection limit of the Qubit assay. DNA protocols using QiaSymphony (DNA_P1 and DNA_P2) achieved the highest median DNA concentrations from pig feces (n=2, 60,2 (range 44,8 - 75,6) ng/µl and 42,1 (range 36,5 - 47,6) ng/µl) of all methods tested (Supplementary Table 1), while DNA_P3 and DNA_P4 resulted in lower and comparable DNA concentrations (Fig 3a).

**Fig 3.**
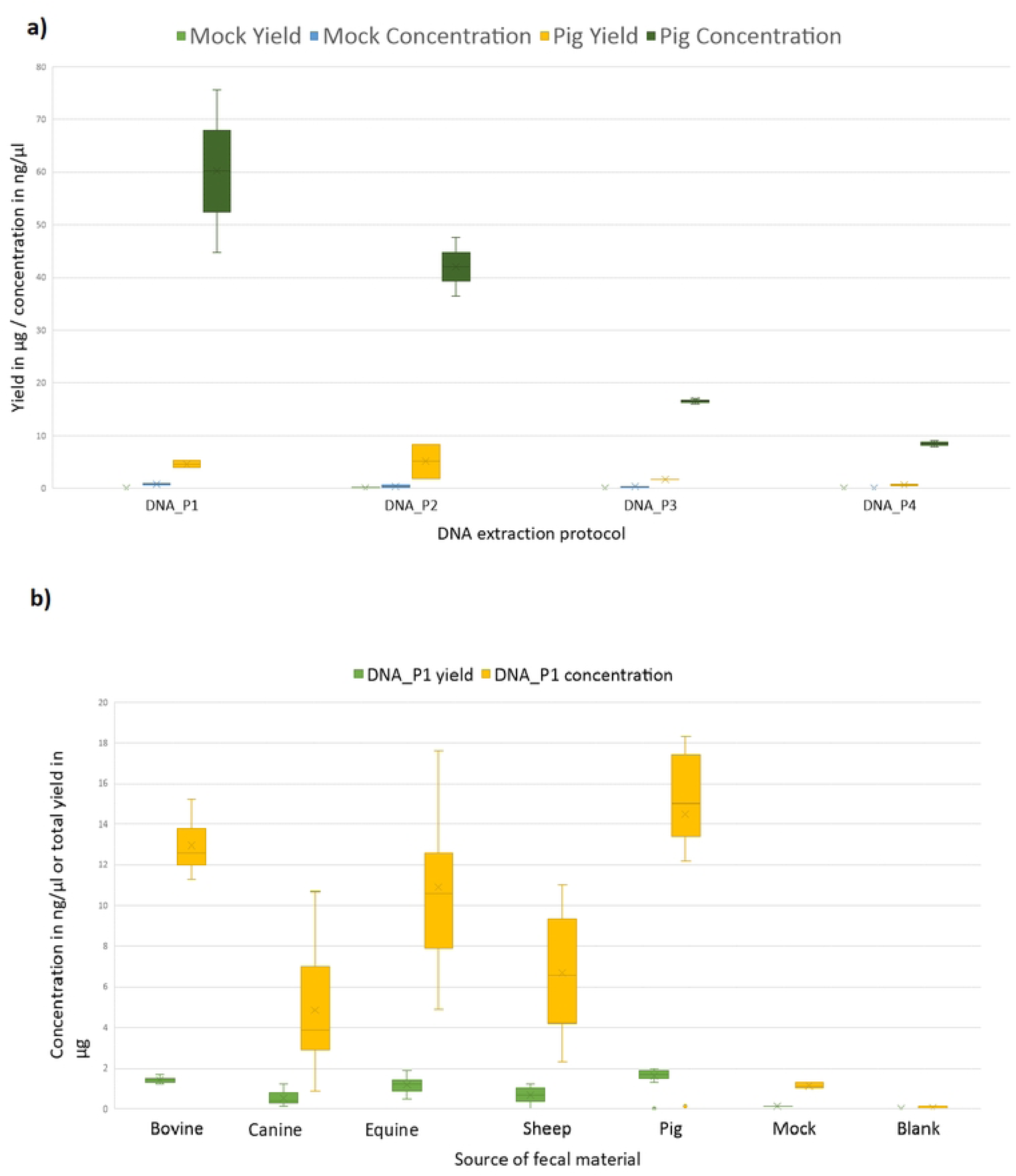
Box and whisker plot depicting the distribution of DNA yields and concentrations achieved using four different extraction protocols for pig, mocks and blanks (a) and pigs, equine, sheep, canines and bovines (b). The central ‘X’ within each box corresponds to the median DNA yield/concentration, while the width of the box represents the interquartile range (IQR) and signifies the variance in DNA yield/concentration across samples. The whiskers extend to show the range of data within 1.5 times the IQR. This plot provides a comprehensive view of the variability in DNA extraction efficiency among the different protocols employed.

### Depiction of the mock communities

The DNA of the mock communities was extracted using protocol DNA_P1 through DNA_P4 and subjected to 16S rRNA amplicon sequencing. The range in the number of classified reads from each mock community was 5 885 (DNA_P4) – 216 557 (DNA_P2), of which 0 – 453 reads were not one of the eight bacterial species in the mock community. The average relative abundance of *L. monocytogenes* was 98.8% (SD 0,25%), which was higher than expected for the mock community (theoretical relative abundance of 95.7%) (Supplementary Fig S1). The average relative abundance of the remainder of the bacteria expected in the mock samples were 0.85% (SD 0.18%) *P. aeruginosa*, 0.32% (SD 0.07%) *B. cereus*, 0.01% (SD 0.01%) *S. enterica* and 0.01% (SD 0.01%). *S. aureus, E. faecalis and L. fermentum* were not detected in any of the mock extracts. In mock extracts created using DNA_P2 (n = 1) and DNA_P3 (n = 1) and DNA_P4 (n=2), none of the sequenced reads matched to *E. coli* nor *Salmonella*.

### Defining a suitable sequencing effort – shotgun metagenomic sequencing

Fecal material on DBS cards from five individuals of each species was purified with the DNA_P1 protocol, using three technical replicates for each individual animal (Fig 3b). The average concentration and total yield are given in Supplementary Table 1.

DNA from 25 fecal samples from the five different animal species, five mocks and five blanks were extracted using DNA_P1 and subjected to shotgun metagenomic sequencing. The average number of reads per fecal sample (irrespective of animal species) was 2,1 x 10^7^ PE reads [SD ± 4,5 x10^6^, range 6,0 x 10^6^ – 2,4 x 10^7^], and the average number of reads for each dataset over species were comparable between bovine, sheep, pig, and equine samples, while the average number of reads for canine samples was slightly lower than for the other species (Table 2, Supplementary Table 2).

**Table 2.** Sequence effort over samples.

| DNA extraction method | Source | Average # PE reads [SD] | Range # PE reads |
| --- | --- | --- | --- |
| DNA_P1 | All animals | $2.1 \times 10^7$ [SD$\pm 4471759$] | $6.2 \times 10^6$, $2.4 \times 10^7$ |
| | Bovine | $2.1 \times 10^7$ [SD$\pm 3708954$] | $1.6 \times 10^7$, $2.4 \times 10^7$ |
| | Canine | $1.7 \times 10^7$ [SD$\pm 6541075$] | $6.2 \times 10^6$, $2.4 \times 10^7$ |
| | Equine | $2.3 \times 10^7$ [SD$\pm 2048548$] | $1.9 \times 10^7$, $2.4 \times 10^7$ |
| | Sheep | $2.1 \times 10^7$ [SD±3494134] | $1.6 \times 10^7$, $2.4 \times 10^7$ |
| | Pig | $2.1 \times 10^7$ [SD±5084972] | $1.2 \times 10^7$, $2.4 \times 10^7$ |

To decide on the appropriate sequencing depth needed to reliably define the functional and taxonomic composition of fecal matter of the five sampled species, the datasets were randomly subsampled at 1 x 10^4^, 5 x 10^4^, 1 x 10^5^, 5 x 10^5^, 1 x 10^6^, 5 x 10^6^ and 1 x 10^7^ reads, creating in total 560 datasets. Each dataset was thereafter taxonomically classified using the TALOS pipeline’s inbuilt Kraken2 with the standard database plus protozoa, fungi & plant (plusPFP) (Standard is: Prokaryotic RefSeq genomes, virus genomes and plasmids). The result of these analyses is presented in Fig 4. The number of observed genera (richness) increased for all datasets when more reads were analyzed, especially between 1 and 5 million reads, albeit the increase in richness tapered rapidly off for blank samples (Fig 4a). After 5 million reads the number of new observed genera per added read decreased (Fig 4a), and the Shannon diversity for all samples was unchanged for datasets larger than 5 million reads (Fig 4b). We did observe larger standard deviations in Shannon diversity for samples from pig and equine samples due to more observed genera for these samples (Fig 4b).

**Fig 4.**
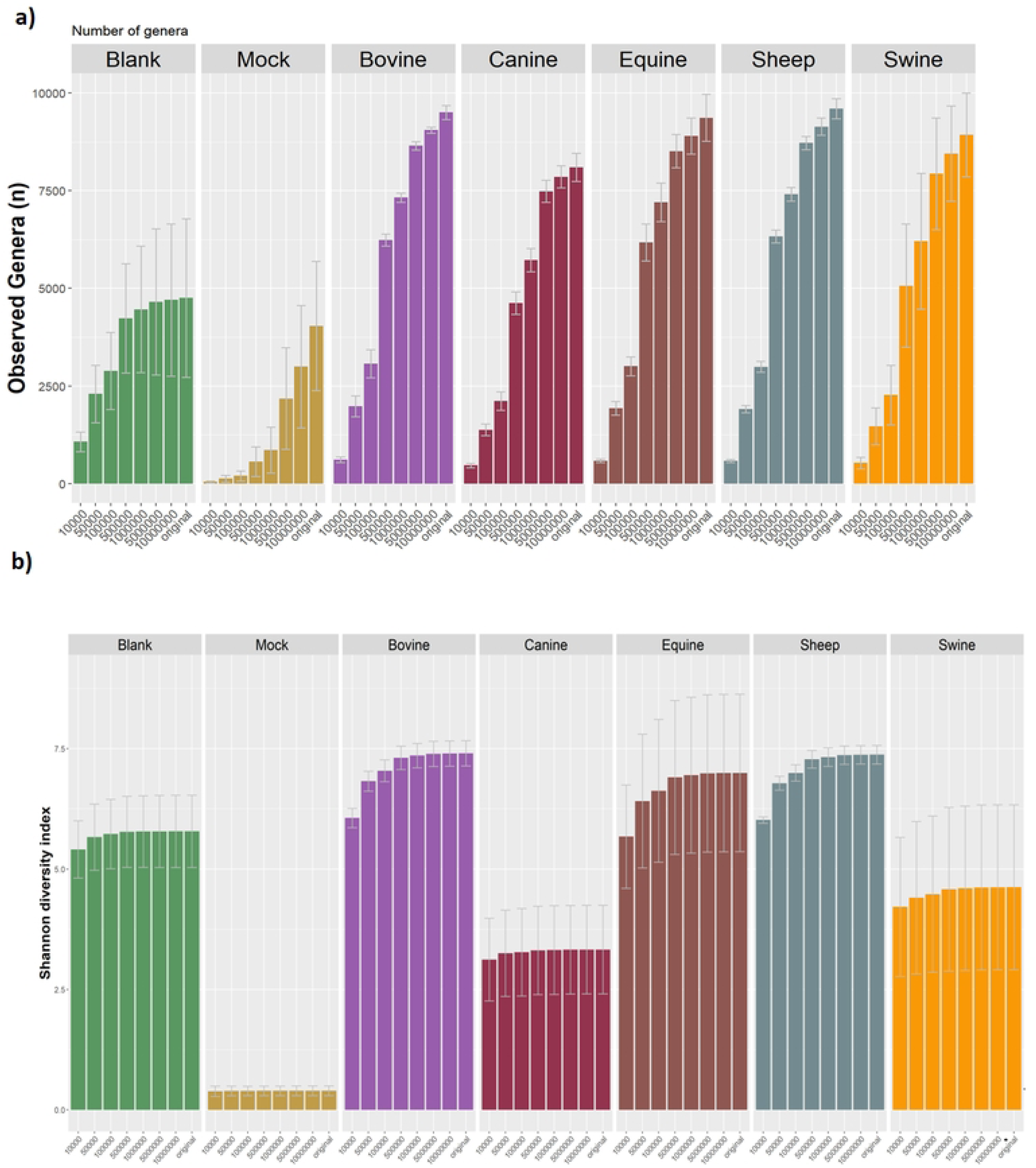
Observed genera (a) and Shannon diversity (b) for all samples in relation to the subsampling depth. Each bar represents the average value for the five samples. Error bars indicate the standard deviation. As little as 5*10^4^ reads was sufficient to capture the Shannon diversity of mock samples, while blank samples Shannon diversity tapered off after 5 x 10^5^ reads. For fecal samples, the Shannon diversity was unchanged after approximately 5 million reads.

Based on the full dataset, the orders Bacteroidales and Eubacteriales (previously Clostridiales) dominate most fecal samples (Fig 5). However, there are species specific differences. For instance, Lactobacillales were more abundant in several of the pig samples. Another difference between the host-species microbiomes is the fraction of classified reads in the microbiomes. For canine fecal microbiomes the classified fraction of the microbiome is much larger than for the bovine microbiomes. Aside from taxonomic classification, DNA sequence composition based on k-mers can be used to cluster microbiome samples with the software sourmash. In this analysis, mock communities clustered together with short branch length due to their great compositional similarity (Fig 6). Animals of the same species and blanks clustered together, with longer branch lengths indicating greater variability between individual animals and blank samples due to random effects (Fig 6).

**Fig 5.**
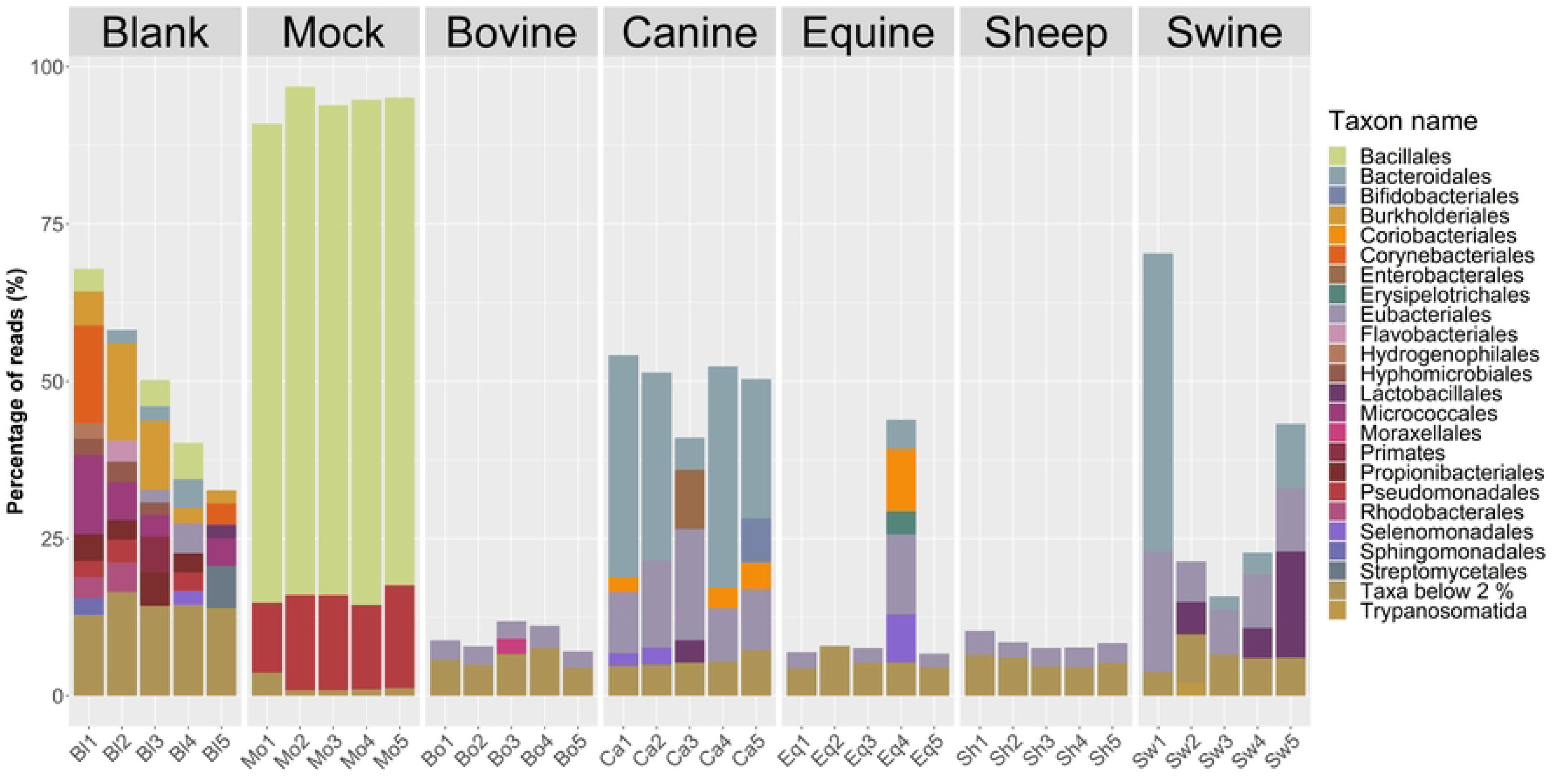
Relative abundance of orders across samples and species for the full dataset before subsampling. A low percentage of reads were classified for sheep, bovine and except for one sample, equine, while for canine and pig feces higher reads classification levels are observed. In total 66 taxa at the order level were identified with a minimum abundance of 0.1%. We only show taxa with a minimum abundance of 2% in at least one sample, all other taxa were group as “Taxa below 2 %” in the legend. Bl: Blank, Mo: Mock, Bo: Bovine, Eq: Equine, Sh: Sheep, Sw: Swine

**Fig 6.**
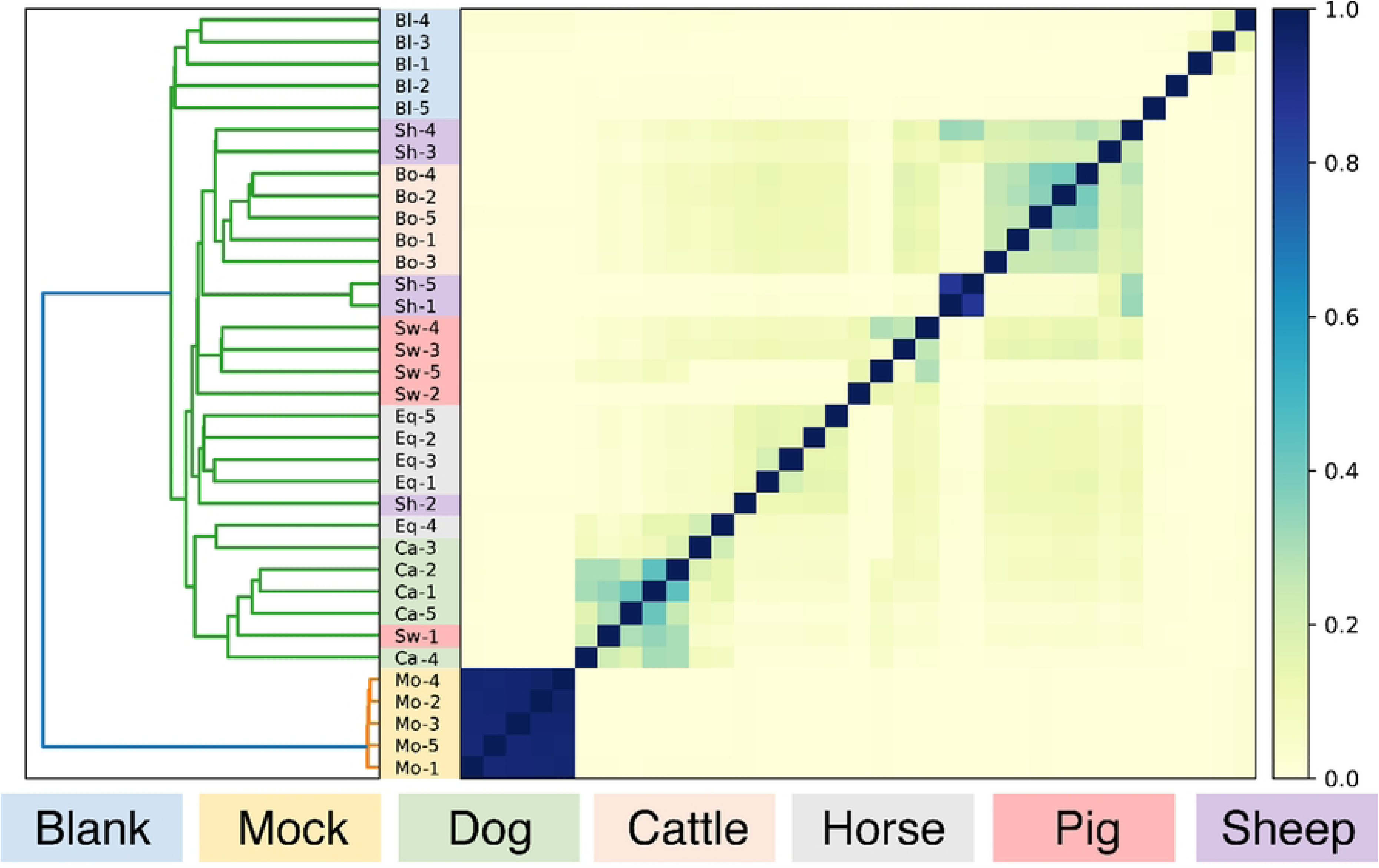
Sourmash cluster analysis of metagenomic datasets of animal’s fecal material, blanks, and mock communities. Sourmash creates a kmer signature for each of the datasets, which than is used with angular similarity as a distance matrix to cluster the samples. Note that the fecal microbiomes of sheep and bovine cluster, and that mock and blank samples are outgroups of the fecal microbiomes.

The functional potential present in the microbial populations was characterized through analyses of the Illumina shotgun metagenome data. The number of reads functionally classified per sample varied depending on the sample type (Supplementary table 2). Blank samples had on average 1.5 million classified reads (sd: ±2.5million), while Mock samples had 12.5 million classified reads (sd: ±1.2 million). The number of classified reads for the animal samples was sheep (4.2 million ± 1.1), bovine (4.8 million ± 0.8), equine (4.9 million ± 1.2), pig (5.7 million ± 1.7) and canine (6.3 million ± 2.1).

The mock and canine samples had the highest percentages of reads matching sequences in the SEED database (57,6% ± 0,6% and 38,2% ± 4,1%, respectively), while the sheep samples had the least reads matching (20,3% ± 3,3%) (Fig 7). The SEED subsystems with most reads assigned regardless of sample type classified to the metabolism of carbohydrates (5,0% ± 2,5%), followed by protein metabolism (3,2% ± 0,5%) and amino acids and derivatives (2,8%±0,9%) (Fig 7). On average, the three most abundant SEED functions for all samples combined were TonB-dependent receptors (1,3% ± 1,4%), β-galactosidase (0,8% ± 0,7%) and multi-antimicrobial extrusion proteins (MATE) (0,7 ± 0,3%). These are functions resembling uptake (TonB-dependent receptors) and export (MATE) functions, while Beta-galactosidase plays an essential role in carbohydrate metabolism. The blank samples had the lowest average number of observed metabolic functions (10.702 ± 4.520), while bovine samples had the highest number of metabolic functions (20.931 ± 1.940).

**Fig 7:**
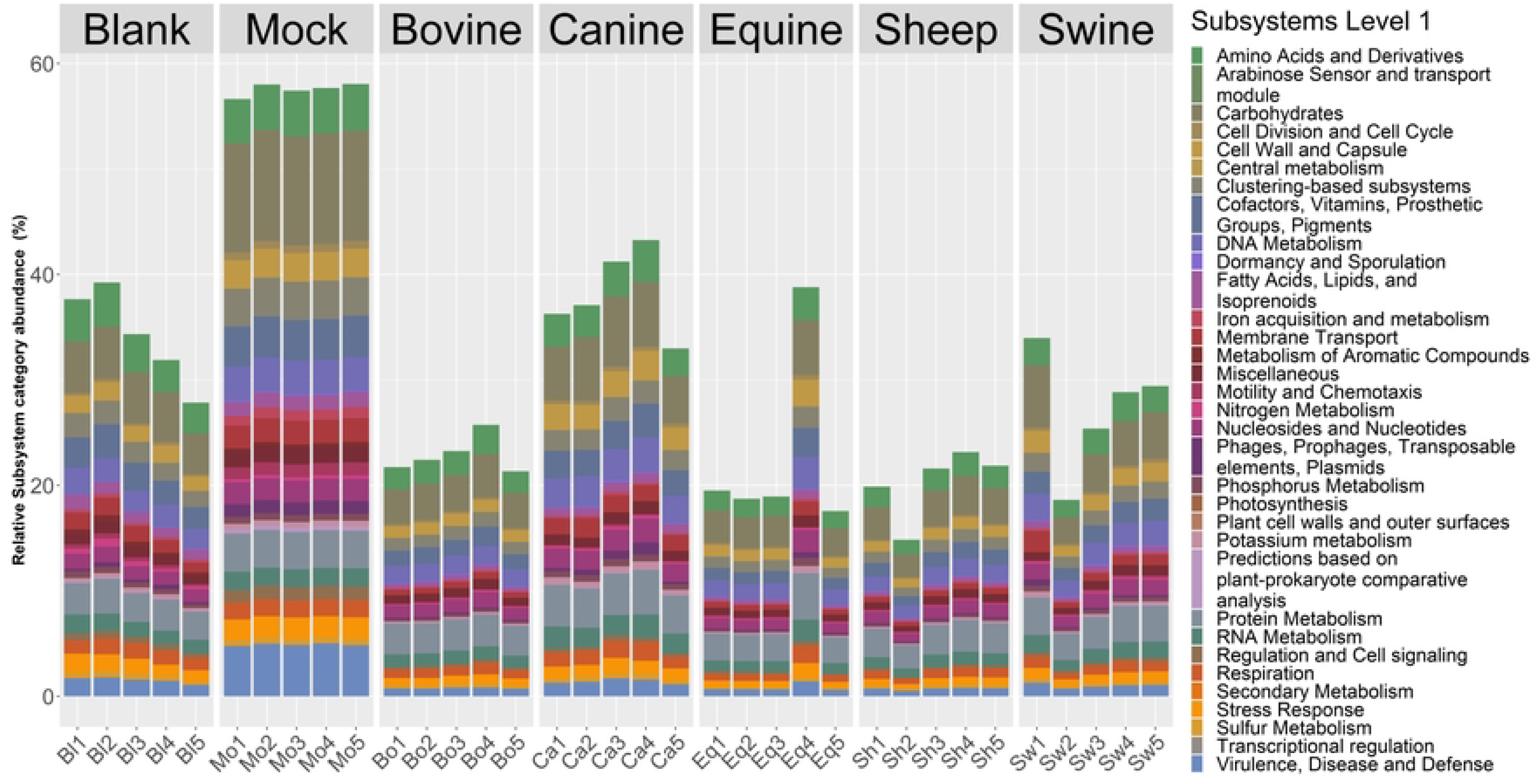
Overview of functional subsystem groups for all samples. Relative abundance of reads classified to different SEED subsystem level 1 categories. Note that Bovine, Sheep and Equine have the lowest levels of classified reads.

Using the subsampled datasets at 1 x 10^4^, 5 x 10^4^, 1 x 10^5^, 5 x 10^5^, 1 x 10^6^, 5 x 10^6^ and 1 x 10^7^ reads, the 560 datasets were subjected to functional classifications. The number of metabolic functions increased with increased sequencing depth and did not taper off for any of the fecal samples nor the mock samples. For the blanks we observed a levelling off in the number of metabolic functions after 5 million reads, due to the limited number of reads produced in these samples (Fig 8). To explore the dynamics of observed functions with sequencing depth in detail, we divided the metabolic functions in two groups 1) infrequent functions, i.e. functions with less than 10 reads, and 2) frequent functions that have 10 or more reads. We then determined for the different subsampling depths the number of infrequent functions (Fig 8). That shows that the number of infrequent functions increases with sequencing depth for the animal fecal microbiomes, while this was not the case for the blank and mock samples. This indicates that the used sequencing depth was sufficient to observe most functions for the blanks and mocks.

**Figure 8:**
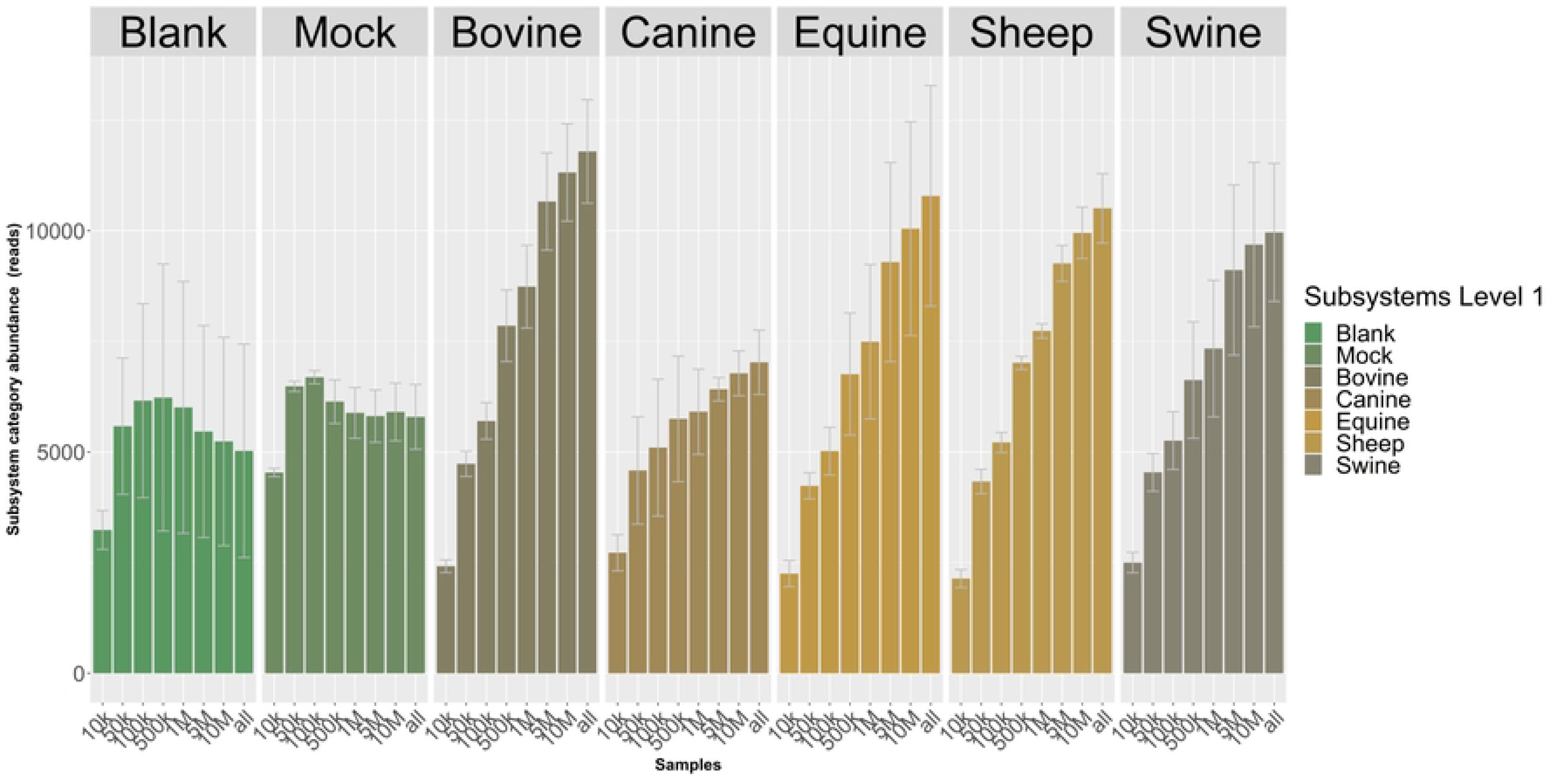
Overview of the number of functions that have less than 10 reads assigned for different sizes of the metagenomic communities. Each bar represents the average value for the five samples. Error bars indicate the standard deviation.

In contrast, the frequent function class increased with increasing sequencing depth for all sample types (data not shown) since functions move from the infrequent group to the frequent group with increasing sequencing depth. These results indicate that our sequencing depth for metabolic functions was not saturated for the animal fecal microbiomes.

One group of metabolic functions that is of broad interest are antimicrobial resistance genes. In the SEED classification system, we find these genes under the subsystem level 2 category: “Resistance to antibiotics and toxic compounds”. Within that category, we find that several classes dominate in the animal fecal microbiomes (Supplementary Fig S1). Those are cobalt-zinc-cadmium resistance, copper homeostasis, multidrug resistance efflux pumps, and resistance to fluoroquinolones. Among the minor classes present in fecal microbiomes are beta-lactamases, enzymes that can breakdown a variety of antibiotics. This class of enzymes had the highest abundances in both the Blank (482 reads per million reads (RPM) ±246) and Mock (431 RPM ± 14) samples. Beta-lactamases were found in the highest levels from canine samples (355 RPM ± 35) and in low levels from sheep feces (150 RPM ± 37) (Supplementary Figure S1).

As we are interested in understanding the detection of AMR genes in our metagenomes, we selected one of the more abundant Beta-Lactamase gene functions and analyzed the abundance with increasing sequencing effort. The function we picked was: “Beta-lactamase_class_C_and_other_penicillin_binding_proteins”. This function could only be detected in the animal fecal microbiomes with 100 reads or more when the sequencing effort was above 1 million reads (Supplementary Figure S2). For other functions, the sequencing effort had to be higher. For example, the function “Regulatory_protein_BlaR1” was only detected in two samples, with more than 100 reads, when using all the reads sequenced. This shows that our sequencing effort was insufficient to detect low abundance AMR genes.

## Discussion

In this study, we identified an efficient, high-throughput DNA extraction method that gives high yield, quality, and unbiased DNA from canine, sheep, equine, bovine, and pig feces spread on DBS cards. Upon shotgun metagenomic sequencing, the resultant DNA successfully determines the taxonomic and functional composition of the metagenome. To boost sensitivity of rare functional traits, increased sequencing effort is needed. This data will be reusable for various scientific studies, hence provide a path towards a more sustainable use of metagenome sequence datasets for One Health scientific discoveries.

Ruminants and horses are fore- and hindgut fermenters, respectively dependent on a well-functioning gut microbiota in the rumen and ceca for the digestion of insoluble grass feeds. The microbiota in turn provides their hosts with nutrients such as short chain fatty acids and proteins (45–47). For monogastric animals, the intestinal microbiota contributes to gastrointestinal health. Severe disturbances to the composition of the gut microbiota (dysbiosis) is associated with progression and development of gut disease such as acute hemorrhagic diarrhea syndrome (8) and colonic cancer (48, 49). Studies on the composition of gut microbiomes of these five animal species are therefore important.

HUNT One Health is the first study that used DBS cards to collect fecal material from canines, sheep, equines, bovines, and pigs intended for metagenomic analysis. As DBS cards lack DNA-stabilizing chemicals, concerns were raised on the suitability of these cards to preserve the fecal samples, or the microbial DNA contained within feces well enough to allow for reliable qualitative and semi-quantitative metagenomic characterization. Especially degradation of genetic material during storage and transport was a concern. The cards were not designed to limit cross-contamination between samples and can only collect a small volume/mass of the samples, possibly reducing the usability of fecal samples for metagenomics. When these challenges are acknowledged during planning, they can be addressed. For instance, the project coordinators provided the owners with specialized pouches with barcodes to protect each card and eliminate cross-contamination. These pouches also contained a small desiccant (silica gel) to help dry the samples. A Takara library preparation specialized for samples of low biomass was used to overcome the challenge of low sample volume. With time, as technology advances, the need for higher DNA amounts for sequencing reaction are also expected to decrease.

DNA extraction can induce various types of bias affecting the inference of the metagenome composition, as shown in several studies. The main goal for this study was therefore to identify a DNA extraction method suitable to extract DNA from 3000 fecal samples collected on DBS cards. The final method had to be suitable for metagenome DNA extraction from feces samples of all five animal species. The efficacy of a DNA extraction method is determined by the amount and type of sample that can be processed properly and the capacities of the extraction kit to cope with chemical and physical inhibitors, including the filter paper itself, the requirements for equipment and training of personnel, etc. (reviewed in (50)). The DBS sample approach is not standardized for amounts of feces on the cards, so irregular amounts of feces will result in some variance in the quantity of input material in the DNA extraction. We tried to limit this by using four discs and trained personnel. An alternative that would increase the throughput time, would be to measure the fecal material prior to processing via gravimetric measurements. A further problem occurs when extracting DNA from fecals from different animal species; incompletely digested dietary material such as fibers in stool differs between individuals and groups such as between herbivores and omnivores (51, 52), resulting in inconsistent amounts of microbes available from the same amounts of feces from different animals. Also, the protein and fatty content of the fecal, as well as more global chemical properties such as pH, vary and can considerably affect the extraction protocol’s overall efficiency. A successful depiction of the microbial community of feces will also be dependent on the community itself, and that varies between species and individuals (53). Ruminants, hind-gut fermenters and monogastric animals exhibit significant differences in their digestive systems. Ruminants possess a multi-chambered stomach, including the reticulum, rumen, omasum, and abomasum, which enables them to digest plant materials through fermentation. Monogastric animals have a simple stomach structure with a single compartment. These different structures will harbor microorganisms that perform digestive tasks on different materials. Therefore, the microbiome differs significantly between species (53, 54). Some of the microbes may be protected from lysing enzymes by encapsulation in various fecal materials. In such cases rigorous crushing could aid in releasing the microbes into solution and lysing. Some spore-forming parasites like *Eimeria*, have though walls preventing DNA from being released into solution unless cells are first rigorously treated. DNA extraction targeting “all” organisms present in stools, would very likely benefit from an efficient homogenizing bead-beating steps (55). Finally, for the HUNT One Health study, the sampling was performed by the animal owners, who had received no prior training and had to follow instructions provided on paper and online video. It is reasonable to assume that they performed the sampling, smearing of samples on faecal cards and successive air-drying variably. Large variations were observable between samples derived from the same animal species, and some cards had obviously not been sufficiently dried before inserting them in the closed pouches for storage and shipping to the HUNT centre (unpublished data). All these variables make quantitative metagenomics, especially for rarer traits, challenging, and this dataset is therefore most suited for qualitative metagenomics.

The mock community in the microbial community standard can be used to assess the quality of extraction, possible bias, and errors in the process. Of the eight microbes from the mock community, all bacteria present at 0.069% or more were identified in one or more of the samples. The relative abundances of the extracted DNA deviated somewhat from the theoretical composition of the mock community, most notably with a slightly higher abundance of *L. monocytogenes* and *B. subtilis* and lower abundance of *P. aerginosa* and *E. coli*, i.e. a small shift towards Gram-positive bacteria. This could indicate a bias in either the extraction method, amplification process, sequencing or bioinformatic pipeline. All four protocols used in this study included a homogenizing step of six minutes bead-beating on the efficient FastPrep instrument. Zhang *et al.* (2021) found that increasing bead-beating time, especially over four towards nine minutes, correlated with a higher abundance of Gram-positives and reduced recovered Gram-negative bacteria (56). Further, nanopore sequencing is known to be error-prone, and as the existing OTU-based approach require at least 97% sequence identity, such sequencing have been considered unsuitable for taxonomic classification of MinION™ reads (34).

With the start of the metagenomics era, the need for the inclusion of control samples arose, followed by discussions on how to deal with sequence data from such controls and the choice of appropriate filter methods to remove sequences that stem from potential contaminants.

Negative (blank) controls have been used in many studies, to subsequently identify and remove taxa present in those controls as potential contaminants, based on the assumption that prevalence of contaminants will naturally be higher in controls than in the samples because of absence of competing DNA (57). However, for this approach to work the input biomass of all sequenced samples must be equal (58). This is a problem for blank samples, which usually contain very minute amounts of DNA. In our hands, sequencing the blanks generated enough reads which were used for community analysis (Supplementary table S2), and the diversity analysis and taxonomic classification indicated a blank community consisting of many different taxa. However, most of these taxa were present in small number of reads only (<50 reads) and the classifications are therefore possibly spurious. However, some of the taxa are most certainly present (e.g. Burkholderiales, Micrococcales), which makes sense as our blanks were not sterile, neithr aseptically handled and included to depict what was in the paper used to collect the fecal samples. Further, the bioinformatic analysis did not apply a lower threshold for reporting taxa, so all classified reads will be reported, regardless of how rare they were, explaining the observed richness of these samples. The community of the blanks do differ remarkably from the fecal samples and the mock communities, demonstrating that neither the collection method nor the kitome influenced the fecal community.

In general, our experiments found several methods to be useful, and we were able to perform an informative decision on which method will work best in the HUNT One Health study. Qiagen’s Blood and Tissue kit performed on the QiaSymphony platform achieved the highest sensitivity and fulfilled our needs for extraction method. In addition, we found 20 million PE reads sufficient to perform robust taxonomic estimates of the metagenomes, as in line with the findings of Hillmann *et al.* (2018)(59). For the functional classification, 20 million PE reads are enough to map out the most common functional pathways, but the method has too low sensitivity to detect the abundance of rarer traits. For instance, we could only detect “Beta-Lactamase Class C and other penicillin binding proteins” in samples with more than 5 million reads, and then only 100 reads assigned to this function. In addition, this function is a “bucket” for many closely related ARG genes, and it thus suggests that for the analysis of specific ARG genes we need more sequencing effort to obtain values that can be used for testing experimental differences. This is also in line with other published studies; for instance, 80 million reads were used to do comparative studies on the abundance and type of AMR genes in pigs and chickens (4). Our dataset is however well fit for qualitative detection, but careful interpretation is necessary as low abundance AMR genes are likely to go undetected. The Norwegian AMR situation is favorable, so low levels of AMR genes are expected in this material.

## Acknowledgements

The analysis of the sequence data was performed under project nn9305k on the SAGA compute cluster provided by Sigma2 - the National Infrastructure for High Performance Computing and Data Storage in Norway. Funding was provided by the Ministry of Agriculture and Food (ref.no. 17/862-7), Regional Forskningsfond Midt-Norge (grant no. 282452), Nord-Trøndelag County authority, Norges Bondelag, Norsk Bonde-og Småbrukarlag, Animalia and the HUNT4-study.

## References

1. Yamamoto EA, Jørgensen TN. Relationships between vitamin D, gut microbiome, and systemic autoimmunity. Front Immunol. 2019;10:3141.

2. Qin J, Li Y, Cai Z, Li S, Zhu J, Zhang F, et al. A metagenome-wide association study of gut microbiota in type 2 diabetes. Nature. 2012;490(7418):55–60.

3. Jaacks LM, Vandevijvere S, Pan A, McGowan CJ, Wallace C, Imamura F, et al. The obesity transition: stages of the global epidemic. Lancet Diabetes Endocrinol. 2019; 7 (3): 231–40.

4. Munk P, Knudsen BE, Lukjancenko O, Duarte ASR, Van Gompel L, Luiken REC, et al. Abundance and diversity of the faecal resistome in slaughter pigs and broilers in nine European countries. Nat Microbiol. 2018;3(8):898–908.

5. Andersen VD, Munk P, de Knegt LV, Jensen MS, Aarestrup FM, Vigre H. Validation of the register-based lifetime antimicrobial usage measurement for finisher batches based on comparison with recorded antimicrobial usage at farm level. Epidemiol Infect. 2018;146(4):515–23.

6. Van Gompel L, Luiken REC, Sarrazin S, Munk P, Knudsen BE, Hansen RB, et al. The antimicrobial resistome in relation to antimicrobial use and biosecurity in pig farming, a metagenome-wide association study in nine European countries. J Antimicrob Chemother. 2019; 74 (4): 865–76.

7. Hendriksen RS, Lukjancenko O, Munk P, Hjelmsø MH, Verani JR, Ng’eno E, et al. Pathogen surveillance in the informal settlement, Kibera, Kenya, using a metagenomics approach. PLoS One. 2019;14(10):e0222531.

8. Herstad KMV, Trosvik P, Haaland AH, Haverkamp THA, de Muinck EJ, Skancke E. Changes in the fecal microbiota in dogs with acute hemorrhagic diarrhea during an outbreak in Norway. J Vet Intern Med. 2021;35(5):2177–86.

9. Herstad KMV, Gajardo K, Bakke AM, Moe L, Ludvigsen J, Rudi K, et al. A diet change from dry food to beef induces reversible changes on the faecal microbiota in healthy, adult client-owned dogs. BMC Vet Res. 2017;13(1):147.

10. Herstad KMV, Vinje H, Skancke E, Næverdal T, Corral F, Llarena AK, et al. Effects of canine-obtained lactic-acid bacteria on the fecal microbiota and inflammatory markers in dogs receiving non-steroidal anti-inflammatory treatment. Animals (Basel). 2022;12(19).

11. Iakhno S, Delogu F, Umu ÖCO, Kjos NP, Håkenåsen IM, Mydland LT, et al. Longitudinal analysis of the faecal microbiome in pigs fed Cyberlindnera jadinii yeast as a protein source during the weanling period followed by a rapeseed- and faba bean-based grower-finisher diet. Anim Microbiome. 2022;4(1):62.

12. Li Y, Gajardo K, Jaramillo-Torres A, Kortner TM, Krogdahl Å. Consistent changes in the intestinal microbiota of Atlantic salmon fed insect meal diets. Anim Microbiome. 2022;4(1):8.

13. Dvergedal H, Sandve SR, Angell IL, Klemetsdal G, Rudi K. Association of gut microbiota with metabolism in juvenile Atlantic salmon. Microbiome. 2020;8(1):160.

14. Fan P, Bian B, Teng L, Nelson CD, Driver J, Elzo MA, Jeong KC. Host genetic effects upon the early gut microbiota in a bovine model with graduated spectrum of genetic variation. ISME J. 2020;14(1):302–17.

15. Holzhausen EA, Nikodemova M, Deblois CL, Barnet JH, Peppard PE, Suen G, Malecki KM. Assessing the impact of storage time on the stability of stool microbiota richness, diversity, and composition. Gut Pathog. 2021;13(1):75.

16. Song SJ, Amir A, Metcalf JL, Amato KR, Xu ZZ, Humphrey G, Knight R. Preservation methods differ in fecal microbiome stability, affecting suitability for field studies. mSystems. 2016;1(3).

17. Tedjo DI, Jonkers DMAE, Savelkoul PH, Masclee AA, van Best N, Pierik MJ, Penders J. The effect of sampling and storage on the fecal microbiota composition in healthy and diseased subjects. PLoS One. 2015;10(5):e0126685.

18. Radhakrishnan ST, Gallagher KI, Mullish BH, Serrano-Contreras JI, Alexander JL, Miguens Blanco J, et al. Rectal swabs as a viable alternative to fecal sampling for the analysis of gut microbiota functionality and composition. Sci Rep. 2023;13(1):493.

19. Li X-m, Shi X, Yao Y, Shen Y-c, Wu X-l, Cai T, et al. Effects of stool sample preservation methods on gut microbiota biodiversity: new original data and systematic review with meta-analysis. mSystems. 2023;11(3):e04297–22.

20. Choo JM, Leong LEX, Rogers GB. Sample storage conditions significantly influence fecal microbiome profiles. Sci Rep. 2015;5(1):16350.

21. Fouhy F, Deane J, Rea MC, O’Sullivan Ó, Ross RP, O’Callaghan G, et al. The effects of freezing on fecal microbiota as determined using MiSeq sequencing and culture-based investigations. PLoS One. 2015;10(3):e0119355.

22. Sinha R, Chen J, Amir A, Vogtmann E, Shi J, Inman KS, et al. Collecting fecal samples for microbiome analyses in epidemiology studies. Cancer Epidemiol Biomarkers Prev. 2016;25(2):407–16.

23. Guthrie R, Susi A. A simple phenylalanine method for detecting phenylketonuria in large populations of newborn infants. Pediatrics. 1963;32(3):338–43.

24. Pribyl AL, Parks DH, Angel NZ, Boyd JA, Hasson AG, Fang L, et al. Critical evaluation of fecal microbiome preservation using metagenomic analysis. ISME Commun. 2021;1(1):14.

25. Taylor M, Wood HM, Halloran SP, Quirke P. Examining the potential use and long-term stability of guaiac fecal occult blood test cards for microbial DNA 16S rRNA sequencing. J Clin Pathol. 2017;70(7):600–6.

26. Albuquerque MCF, Herwaarden Yv, Kortman GAM, Dutilh BE, Bisseling T, Boleij A. Preservation of bacterial DNA in 10-year-old guaiac FOBT cards and FIT tubes. J Clin Pathol. 2017;70(11):994–6.

27. Vandeputte D, Tito RY, Vanleeuwen R, Falony G, Raes J. Practical considerations for large-scale gut microbiome studies. FEMS Microbiol Rev. 2017;41(Suppl_1):S154–67.

28. Hale VL, Tan CL, Knight R, Amato KR. Effect of preservation method on spider monkey (Ateles geoffroyi) fecal microbiota over 8 weeks. J Microbiol Methods. 2015;113:16–26.

29. Skeen HR, Willard DE, Jones AW, Winger BM, Gyllenhaal EF, Tsuru BR, et al. Intestinal microbiota of Nearctic-Neotropical migratory birds vary more over seasons and years than between host species. Mol Ecol. 2023;32(12):3290–307.

30. Bolt Botnen A, Bjørnsen MB, Alberdi A, Gilbert MTP, Aizpurua O. A simplified protocol for DNA extraction from FTA cards for fecal microbiome studies. Heliyon. 2023;9(1):e12861.

31. Sciences NUoL. HUNT One Health

32. Hirai M, Nishi S, Tsuda M, Sunamura M, Takaki Y, Nunoura T. Library Construction from Subnanogram DNA for Pelagic Sea Water and Deep-Sea Sediments. Microbes and Environments. 2017;32(4):336–43.

33. Forcina G, Pérez-Pardal L, Carvalheira J, Beja-Pereira A. Gut Microbiome Studies in Livestock: Achievements, Challenges, and Perspectives. Animals. 2022;12(23):3375.

34. Cuscó A, Catozzi C, Viñes J, Sanchez A, Francino O. Microbiota profiling with long amplicons using Nanopore sequencing: full-length 16S rRNA gene and the 16S-ITS-23S of the rrn operon. F1000Res. 2018;7:1755.

35. De Coster W, D’Hert S, Schultz DT, Cruts M, Van Broeckhoven C. NanoPack: visualizing and processing long-read sequencing data. Bioinformatics. 2018;34(15):2666–9.

36. Curry KD, Wang Q, Nute MG, Tyshaieva A, Reeves E, Soriano S, et al. Emu: species-level microbial community profiling of full-length 16S rRNA Oxford Nanopore sequencing data. Nature Methods. 2022;19(7):845–53.

37. Barnett DJM, Arts ICW, Penders J. microViz: an R package for microbiome data visualization and statistics. J Open Source Softw. 2021;6:3201.

38. Chen Y, Chen Y, Shi C, Huang Z, Zhang Y, Li S, et al. SOAPnuke: a MapReduce acceleration-supported software for integrated quality control and preprocessing of high-throughput sequencing data. GigaScience. 2018;7(1):1–6.

39. Bushnell B. Masked version of hG19 by Brian Bushnell. In: bbmap, editor. Zenodo. 2018.

40. Wood DE, Lu J, Langmead B. Improved metagenomic analysis with Kraken 2. Genome Biology. 2019;20(1):257.

41. Silva GGZ, Green KT, Dutilh BE, Edwards RA. SUPER-FOCUS: a tool for agile functional analysis of shotgun metagenomic data. Bioinformatics. 2015;32(3):354–61.

42. Buchfink B, Xie C, Huson DH. Fast and sensitive protein alignment using DIAMOND. Nature Methods. 2015;12(1):59–60.

43. Pierce NT, Irber L, Reiter T, Brooks P, Brown CT. Large-scale sequence comparisons with sourmash. F1000Research. 2019;8:1006.

44. Oksanen J, Blanchet FG, Kindt R, Legendre P, Minchin PR, O’Hara RB, et al. vegan: Community Ecology Package. Github. 2023.

45. Russell JB. Rumen microbiology and its role in ruminant nutrition. Department of Microbiology, Cornell University; 2002.

46. Mackie RI. Mutualistic Fermentative Digestion in the Gastrointestinal Tract: Diversity and Evolution. Integrative and Comparative Biology. 2002;42(2):319–26.

47. Yen J-T. Digesta processing and fermentation. Encyclopedia of Animal Science. 2004:282.

48. Tilg H, Adolph TE, Gerner RR, Moschen AR. The Intestinal Microbiota in Colorectal Cancer. Cancer Cell. 2018;33(6):954–64.

49. Arnesen H, Hitch TCA, Steppeler C, Müller MHB, Knutsen LE, Gunnes G, et al. Naturalizing laboratory mice by housing in a farmyard-type habitat confers protection against colorectal carcinogenesis. Gut Microbes. 2021;13(1):1993581.

50. Demirev PA. Dried Blood Spots: Analysis and Applications. Analytical Chemistry. 2013;85(2):779–89.

51. Müller CE. Long-stemmed vs. cut haylage in bales—Effects on fermentation, aerobic storage stability, equine eating behaviour and characteristics of equine faeces. Animal Feed Science and Technology. 2009;152(3):307–21.

52. Jackson MI, Jewell DE. Balance of saccharolysis and proteolysis underpins improvements in stool quality induced by adding a fiber bundle containing bound polyphenols to either hydrolyzed meat or grain-rich foods. Gut Microbes. 2019;10(3):298–320.

53. de Jonge N, Carlsen B, Christensen MH, Pertoldi C, Nielsen JL. The Gut Microbiome of 54 Mammalian Species. Animals. 2022;13.

54. Mallott EK, Amato KR. Host specificity of the gut microbiome. Nature Reviews Microbiology. 2021;19(10):639–53.

55. Combrink L, Humphreys IR, Washburn Q, Arnold HK, Stagaman K, Kasschau KD, et al. Best practice for wildlife gut microbiome research: A comprehensive review of methodology for 16S rRNA gene investigations. Frontiers in Microbiology. 2023;14.

56. Zhang B, Brock M, Arana C, Dende C, van Oers NS, Hooper LV, et al. Impact of Bead-Beating Intensity on the Genus- and Species-Level Characterization of the Gut Microbiome Using Amplicon and Complete 16S rRNA Gene Sequencing. Front Cell Infect Microbiol. 2021;11.

57. Davis NM, Proctor DM, Holmes SP, Relman DA, Callahan BJ. Simple statistical identification and removal of contaminant sequences in marker-gene and metagenomics data. Microbiome. 2018;6(1):226.

58. Du J, Zhang J, Zhang D, Zhou Y, Wu P, Ding W, et al. Background Filtering of Clinical Metagenomic Sequencing with a Library Concentration-Normalized Model. Microbiol Spectr. 2022;10(5):e0177922.

59. Hillmann B, Al-Ghalith GA, Shields-Cutler RR, Zhu Q, Gohl DM, Beckman KB, et al. Evaluating the Information Content of Shallow Shotgun Metagenomics. mSystems. 2018;3(6):10.1128/msystems.00069-18.

